# Transitions in human gut viral communities from ancient to modern societies

**DOI:** 10.1101/2025.09.16.676446

**Authors:** Aydin Loid Karatas, Michael Iter, Jacob Kelman, Natalia Shahwan, Sarah Bonver, Natalie A. Falta, Molly M. Fox, Cali Fujimoto, Eleanor Gorham, Yewon Jang, Meena Khan, Madelaine Leitman, Natalie Martin, Robert Morris, Bobbie Patton, Sofia Potter, Amit Rand, Madeleine Swope, Isha Tripathi, Ben Knowles

## Abstract

The composition and function of the human gut microbial community (the microbiome) have changed substantially over millennia, with implications for human health. While microbiome research has focused primarily on bacterial dynamics, the long-term history of gut viral communities (the virome) remains largely unexplored, despite their crucial role in shaping bacterial populations.

We analyzed gut viromes from 14 pre-modern human coprolites (1301 BCE–1400s CE), as well as 502 non-industrialized and 492 industrialized contemporary human fecal samples. We found that, from pre-modern to contemporary and industrialized populations, human gut viral communities have become more similar in gene content, increasingly dominated by temperate lifestyles, and more supportive of bacterial pathogenicity. These synergistic ecological shifts suggest that long-term changes, especially with industrialization, have fundamentally altered the gut virome, likely affecting human health. These insights into historical shifts in gut viral community and function open potential avenues for ecologically grounded therapeutics to enhance gut microbiome resilience.

## Introduction

The composition and function of the human gut microbiome have changed significantly throughout human history, affecting gut function and health^1–3^. However, while there is a growing understanding of the long-term trajectories of bacterial gut communities^4–6^, and although recent work has shown taxonomic conservation and change in viral communities over deep time^7^, it remains largely unknown how viral communities have functionally changed from pre-modern to modern times, especially with industrialization^8,9^.

The gut microbial community assists in digestion, metabolism, and immune response, influencing nearly every aspect of human physiology^10–12^. Imbalances in gut bacterial composition and function, known as dysbiosis, have been linked to a wide range of diseases affecting numerous organs^4,13,14^. Dysbiosis has been extensively studied in both chronic and acute human illnesses, particularly in industrialized societies^3,15^. These studies highlight the roles of diet, sedentary lifestyles, stress, healthcare practices, globalization, and sanitation in driving human illness via disruption of the gut microbiome^3,12,16,17^.

While much research has focused on bacteria, the viruses that infect them (i.e., bacteriophages) are also essential components of the gut microbiome ^8,9,18^. These viruses regulate bacterial community composition and function, playing key roles in human health^19–21^. Bacteriophages can pursue either lytic or temperate infection strategies^22–25^. In lytic infections, viruses rapidly destroy their bacterial hosts. In contrast, temperate infections delay lysis, and may provide immunity from further infection, and promote sustained horizontal gene transfer of functions like pathogenicity^23–25^. Altogether, the distinct viral lifestyles can have divergent and sometimes reinforcing effects on gut microbiome structure, stability, and function, ultimately influencing human health^21,26^.

However, our knowledge of current changes in the gut virome lacks historical context, as the trajectory of the human gut virome from pre-modern to modern times remains unknown^7^. While acknowledging the well-established caveats of ancient DNA-based (aDNA) analyses and their limited sample sizes^27,28^, we used the unique power of ancient DNA to directly observe the human gut virome before the advent of industrialization and the modern era (circa 1500 CE)^7,29^. We analyzed 14 pre-modern fecal gut metagenomes from hunter-gatherer and small-scale, non-industrialized societies across four continents (modern-day Austria, Mexico, South Africa, and the United States; **Table 1**), dating from 1301 BCE to 1400s CE^6,30,31^. We also analyzed contemporary samples for comparison: 492 metagenomes from industrialized societies^6,32^ and 502 from non-industrialized small-scale societies^6,33^.

Using metagenomic and genomic analyses, including the reconstruction of assembled phage genomes (APGs; assembled contigs identified as viral and quality-filtered with CheckV; n = 42, n = 2316, n = 903 pre-modern, non-industrialized, and industrialized APGs), we investigated shifts in viral functional composition, lifestyle, and pathogenicity. Compared to pre-modern coprolite samples, contemporary samples—especially those from industrialized societies—contained a higher proportion of temperate viruses and exhibited elevated gene-content similarity. Additionally, we observed that viromes from industrialized societies harbored elevated frequencies of pathogenicity genes. Altogether, our results suggest that viruses have played a major role in reshaping the human gut microbiome, potentially contributing to the increased prevalence of gut dysbiosis and chronic disease observed in industrialized societies^14,17,19^. Understanding these long-term virome shifts may help guide ecologically founded therapeutics for restoring gut function and resilience in modern populations.

## Methods

### Sample aggregation

Summaries of all samples are shown in **Supplementary Table 1**, including extended details of all samples, their sources, and bioinformatic metrics.

We aggregated previously published current-day industrialized and non-industrialized samples (industrialized samples from China, Denmark. Spain and USA; non-industrialized samples from Fiji, Madagascar, Mexico, Nepal, Peru, and Tanzania; samples and categorization as industrialized or non-industrialized from 6) with pre-modern samples spanning the Bronze Age at 1301 Before Current Era (BCE) to 1400s Current Era (CE) from disparate sites in Austria, Mexico and the United States (Zape, BMS, and AWC sites) and South Africa. Altogether this comprised 492 samples from industrialized populations (China, n = 74^32^; Denmark, n = 109^6^; Spain, n = 140^6^; USA, n = 169^6^), 502 samples from non-industrialized populations (Fiji, n = 174^6^; Madagascar, n =112^6^; Mexico, n = 22^6^; Nepal, n = 56^33^; Peru, n = 36^6^; Tanzania, n = 102^6,33^), and 14 pre-modern samples (Austria, n = 3^31^; AWC, n = 36; BMS, n = 26; Zape, n = 3^6^; South Africa, n = 3^30^).

### Sample downloading and fastq generation

A work flow summary is shown in **Supplementary Figure 1**. All bioinformatic code is available at https://github.com/hopefulmonstersucla/coprolite_viromes.

Publicly available metagenomic sequencing datasets were downloaded from the NCBI Sequence Read Archive (SRA) using SRA Toolkit v3.0.5 and converted to FASTQ format for downstream analyses. Both single-end and paired-end sequencing libraries were accepted.

### Read quality control

Sequencing reads were quality filtered and adapter trimmed using Trim Galore v0.6.10 under default parameters (https://github.com/FelixKrueger/TrimGalore). Human DNA contamination was then removed by aligning reads to the human reference genome (GRCh38) using Bowtie2 v2.5.2 and discarding aligned reads^34^. Metadata describing read lengths, sequencing depth, and sequence retention during processing were extracted using Seqtk v1.4-r130-dirty (see **Supplementary Table 1**).

### Confirming human gut origin of ancient microbiome samples

To determine if ancient samples originated from the human gut, as opposed to being dominated by sequences from the surrounding soil they were sampled from, MetaPhlAn v4.2.2 was used to generate the species composition of all pre-modern samples, as well as a subset of 24 industrial, 24 non-industrial, and 24 diverse soil samples from across various biomes in Europe and Asia (**Supplementary Table 1**)^6,35^. MetaPhlAn abundance tables of each sample were generated under default settings. Tables were then merged and converted to biom format for analysis with SourceTracker2 v2.0.1 (**Supplement Figure 2**).

Samples that qualitatively matched prior SourceTracker2 results were kept for analysis^6,30,36^.

### Assembly

Quality-controlled reads were assembled into contigs using MEGAHIT v1.2.9 under default parameters^37^. These assemblies (“All contigs”; **Supplementary Figure 1**) formed the basis of all subsequent metagenomic and genomic analyses.

### Confirming ancient sequence origin with pydamage

To confirm ancient sequence origin, samples that passed SourceTracker filtering were further evaluated using PyDamage v1.0^38^. Quality-filtered reads were aligned to sample-specific assemblies using Bowtie2 v2.5.2, and contigs were screened for patterns of ancient DNA damage under default PyDamage parameters. To generate a conservative set of authenticated ancient contigs, only contigs exceeding PyDamage’s default damage threshold of 0.5 were retained for downstream analyses.

### Isolating phage contigs and genomes

We isolated a conservative pool of phage contigs from the assembled contigs with PhaMER^39^ using default parameters except that we increased the phage classification threshold from the developers’ default of 0.3 to 0.5 to provide a more conservative, higher-confidence set of phage contigs (“Phage contigs”; **Supplementary Figure 1**)^39^. This single conservatively defined phage contig pool formed the basis of all downstream metagenomic analyses.

We then extracted complete phage genomes from the “Phage contigs” pool with CheckV v1.0.1 using database v1.5^40^. Accepting only genomes marked as “complete” (checkv_quality = complete) and without contamination (contamination = 0), the most conservative and highest degree of confidence in genome completeness, we identified n = 42, n = 2316, n = 903 pre-modern, non-industrialized, and industrialized genomes, respectively. These genomes made up the Assembled Phage Genome (APG) pool for subsequent genomic analyses (**Supplementary Figure 1**).

### Phage genome protein-sharing network

We implemented a protein-sharing network to investigate the functional relatedness between viruses we had recovered genomes for. To do this, we first used Prodigal within the PROKKA package to call APG ORFs^41,42^ (**Supplementary Figure 1**). The ORFs were then used to make protein clusters with vConTACT2^43^ (v0.11.3, db=ProkaryoticViralRefSeq211-Merged), which were then compared between phage genomes to facilitate clustering (scripts available at https://github.com/hopefulmonstersucla/coprolite_viromes)^44^. Importantly, the RefSeq database served as a broad viral reference, allowing the human gut viromes to be placed within the wider landscape of currently characterized viral genomic functional diversity. The resulting network, characterized by phage genomes as nodes and edge lengths based on the similarity of the proteins they code for, was then visualized in Cytoscape^45^ (v3.10.2; analysis shown in **Figure 2a**; see below for details on permutation analysis).

**Figure 1:**
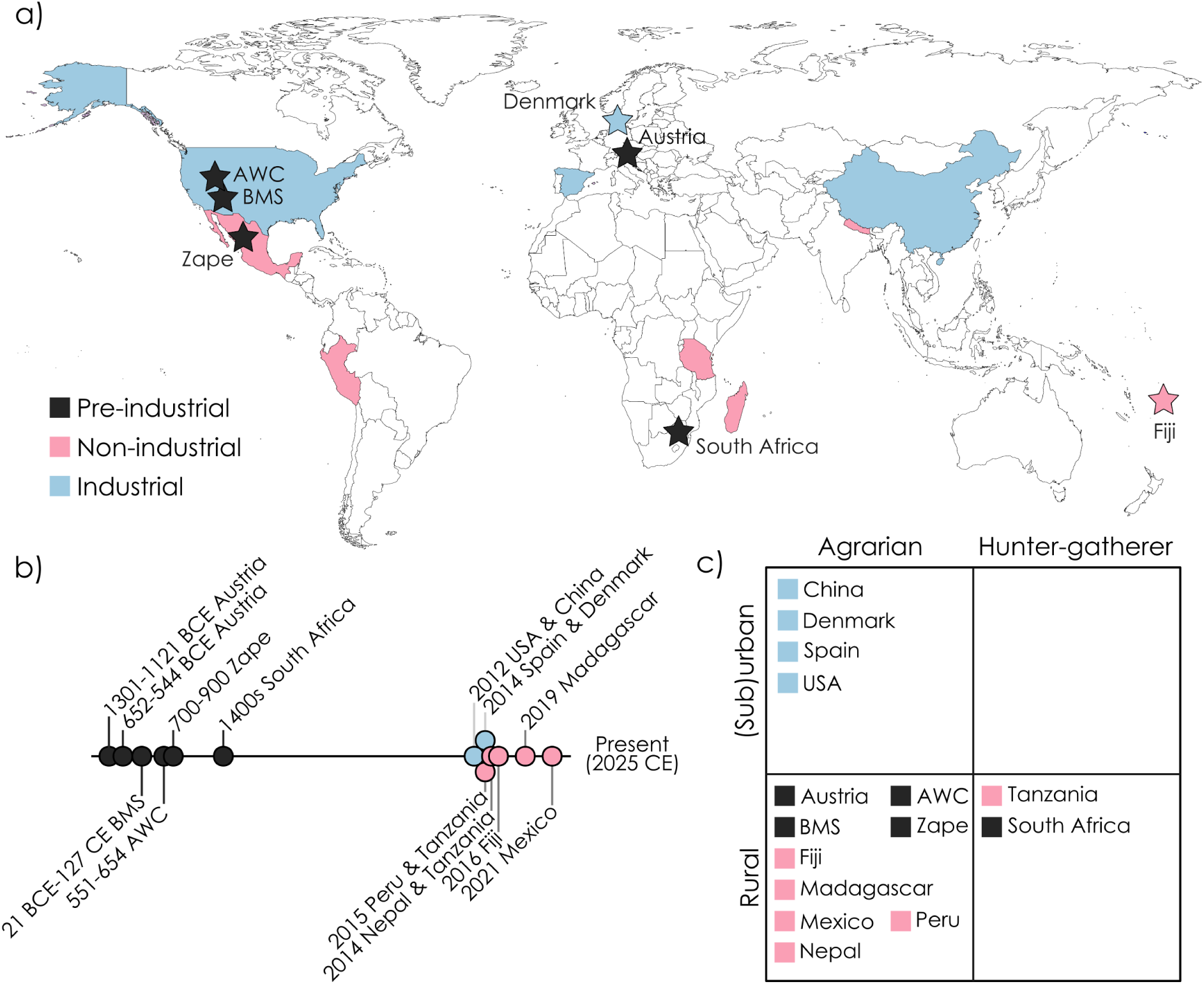
The geographic and temporal origins of pre-modern, non-industrialized, and industrialized fecal samples and their associated human lifestyles. **(a)** Map, **(b)** timeline, and **(c)** social structure and lifestyle of human sources pre-modern fossilized (black, n = 14; coprolites) and non-industrialized (pink, n = 502) and industrialized (blue, n = 492) fresh fecal samples. Note that non-industrialized and industrialized samples are both contemporary. Stars in **(a)** represent point locations or small countries in pre-modern and contemporary samples, respectively. Time points in **(b)** are shown as log_10_-transformed years before present. Years Before Common Era (BCE) and Common Era (CE) values in **(b)** indicate when samples were taken in contemporary samples and are published estimates for pre-modern sample origin, including ranges where appropriate.

**Figure 2:**
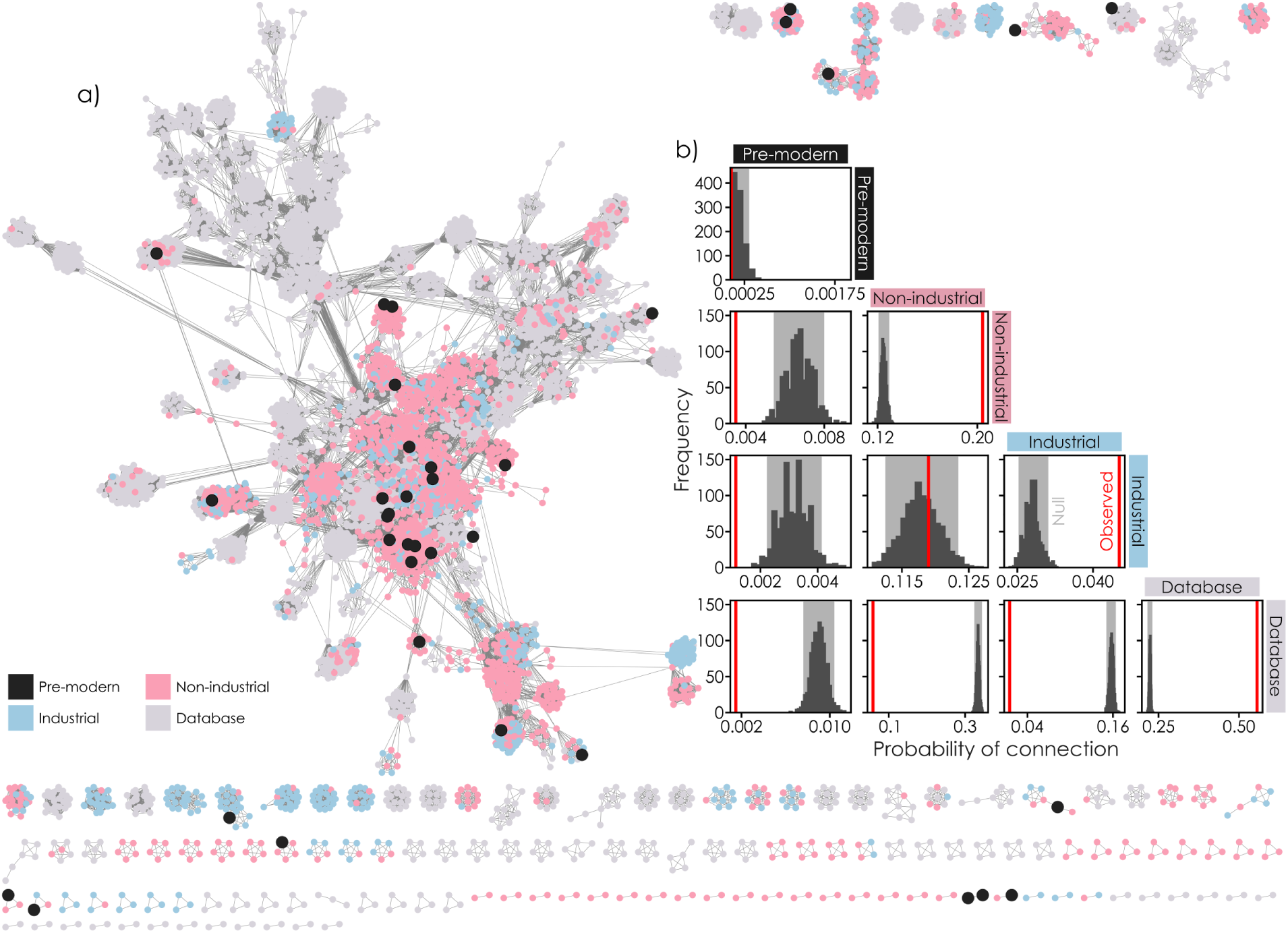
Increasing gene-content similarity in modern vs. pre-modern viruses. **(a)** Shared-protein network of Assembled Phage Genomes from pre-modern (black), non-industrialized (pink), and industrialized (blue) samples and previously characterized phage genomes from the ProkaryoticViralRefSeq211-Merged database (grey; n = 42, n = 2316, n = 903, n = 4534 pre-modern, non-industrialized, industrialized, and database genomes, respectively)^39^. Individual phage genomes are shown as circles (nodes) and connections are shown as grey lines (edges) in the network. **(b)** The frequency of connection (shared edges) between phage genomes in and between each category. Grey histograms show the distribution of connections under random chance generated by permutation with 1000 iterations with shaded areas showing 95 % confidence intervals (CI) and red vertical lines showing observed values. Observed values falling within the 95% CI indicate within-group connectivity similar to random expectation.

### Statistical analysis of protein-sharing network

We contrasted the observed number of connections (i.e., relatedness) between sample categories in the APG protein-sharing network with connections expected by chance (**Figure 2b**). To do so, we used a permutation over 1000 iterations to randomly ‘draw’ two words from a list made up of n = 42 ‘pre-modern’, n = 2316 ‘non-industrial’, n = 903 ‘industrial’, and n = 4534 ‘database’ (i.e., the number of genomes in each category of the network). This was used to generate a null distribution of connections as driven by chance. 95 % CIs of this distribution were defined as excluding the top and bottom 2.5 % of values. Observed connection frequencies falling within the 95 % CI show a lack of similarity, observed values above the 95 % CI show that genomes from within a comparison are more similar than dictated by chance (i.e., they cluster), and observed values falling below the 95 % CI show clustering away from each other.

### Metagenomic viral lifestyle prediction

We quantified the lifestyle of phages in the “Phage contigs” pool using PhaTYP with default parameters^46^. This categorized each unique, non-redundant phage contig as either temperate, lytic, or unpredicted (analysis shown in **Figure 3a**). Python scripts were used to extract the number of contigs in each category in each sample (https://github.com/hopefulmonstersucla/coprolite_viromes).

**Figure 3:**
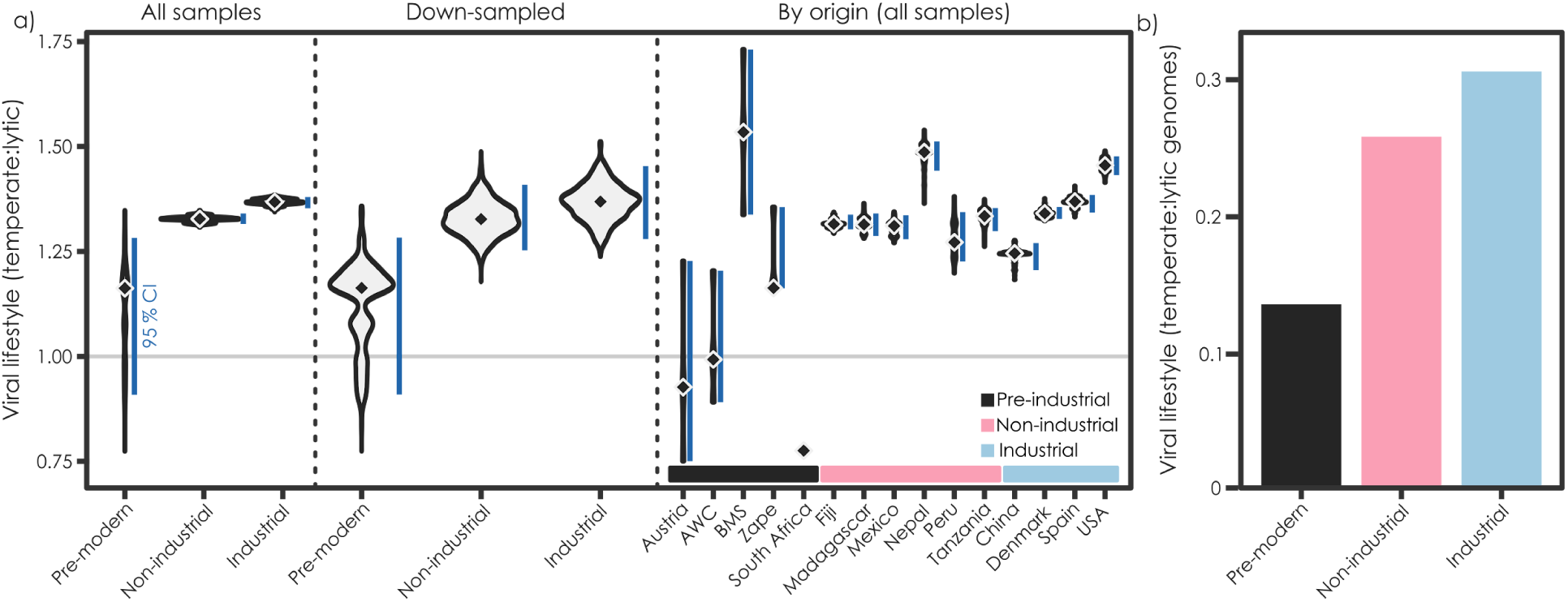
Increased temperateness in modern vs. pre-modern viruses. **(a)** Temperate to lytic ratio of viruses in metagenomically-assembled phage contigs across time periods analyzed using PhaTYP^35^ when viewed as all samples combined, downsampled to account for the smaller pre-modern sample size, and by source location. Bootstrapped value distributions (violins), median values (diamonds), and 95 % confidence intervals (blue vertical lines) are shown from 1000 iterations (n = 14, n = 502, n = 492 for pre-modern, non-industrialized, and industrialized samples, respectively). Values > 1 reflect temperate dominance of the viral community, and < 1 show lytic dominance. **(b)** The prevalence of temperate and lytic assembled phage genomes (APGs) in pre-modern, non-industrial and industrial modern samples (black, pink, and light blue, respectively) as the temperate:lytic genome ratio after panel (**a**) as identified using CheckV^37^ (n = 42, n = 2316, n = 903 pre-modern, non-industrialized, and industrialized APGs, respectively).

Analysis of temperate-to-virulent ratio of phages was limited to samples with at least 100 contigs annotated with either lifestyle. This removed one sample each from Spain (Industrialized), Fiji (Non-industrialized), and South Africa (Pre-modern) due to lack of contigs (samples contained 1, 34, and 5 contigs respectively compared with between 174 and 73701 for all other samples; **Figure 3**).

### Genomic viral lifestyle prediction

In addition to identifying complete phage genomes, CheckV^40^ also annotates whether the genomes are proviral or not (provirus = Yes or No). To complement our metagenomic lifestyle assessment via an independent genomics-based approach, we created a ratio of provirus = Yes vs. provirus = No genomes from the CheckV output to assess the lifestyle of each sample (analysis shown in **Figure 3b**).

### Metagenomic pathogenicity gene frequency

We used PROKKA with the argument “– kingdom Bacteria” to extract ORFs from all phage contigs (“Identify metabolic ORFs” in **Supplementary Figure 1**)^41^. Virulence factors (i.e., pathogenicity genes) were then identified by hidden Markov model (HMM) alignment against the Virulence Factor Database (VFDB) to quantify both the frequency and identity of pathogenicity genes across samples (**Supplementary Figure 1**; analysis shown in **Figures 4a-b**)^47^. VFDB proteins were grouped by Virulence Factor ID and converted into HMM profiles using MAFFT v7.520 and HMMER v3.4^47,48^. Phage ORFs were then queried against these profiles using hmmsearch with a stringent inclusion threshold of “– incE 1e-10”. We filtered for alignments with Full Sequence Scores of at least 25 that were also at least 10-fold higher than the Full Sequence Bias. We quantified the proportion of annotated phage ORFs assigned to virulence factors, together with the identities of detected virulence genes.

**Figure 4:**
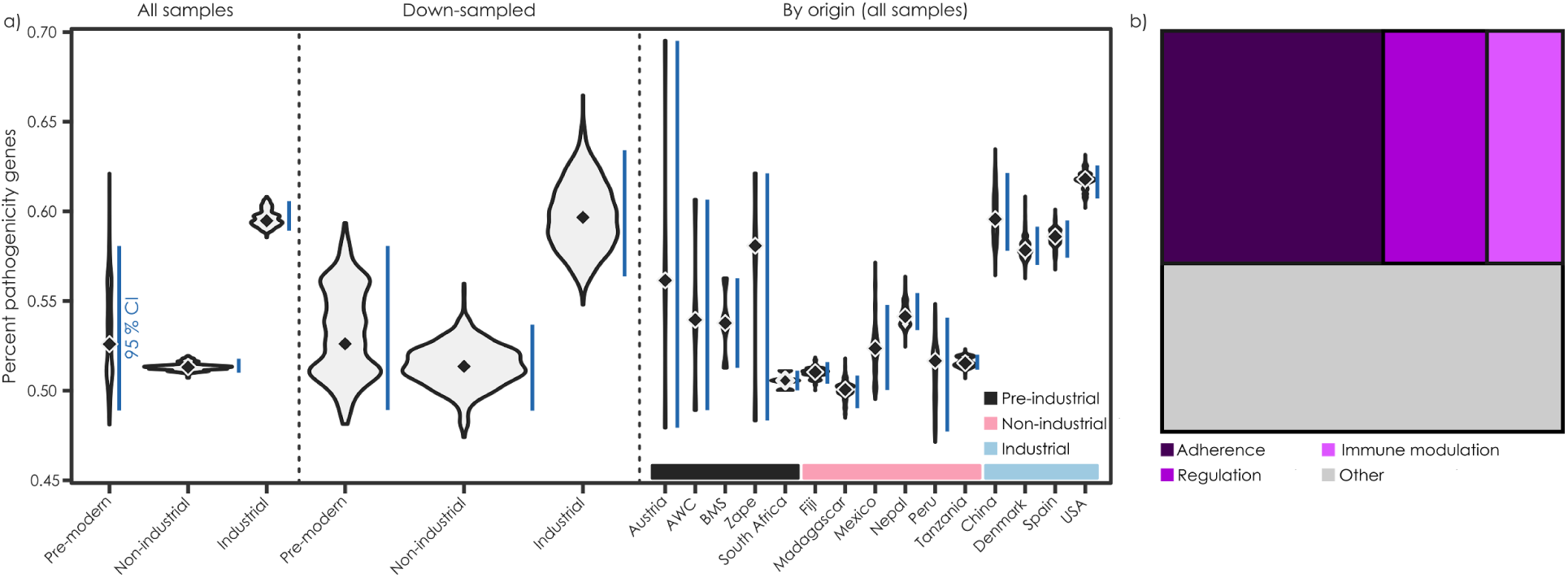
Increased pathogenicity gene frequencies and immuno-regulatory focus in contemporary industrialized viromes vs modern non-industrialized and pre-modern viral communities. **(a)** The percentage of annotated phage ORFs assigned to virulence factors (i.e., pathogenicity genes) in pre-modern, non-industrialized, and industrialized samples when viewed as all samples combined, downsampled to account for the smaller pre-modern sample size, and by source location. Bootstrapped value distributions (violins), median values (diamonds), and 95 % confidence intervals (blue vertical lines) are shown from 1000 iterations. **(b)** The frequency of virulence factors (pathogenicity genes) responsible for adherence, regulation, and immune modulation vs. all other categories in the top 100 most differentially represented virulence factors across samples.

### Statistical analyses of viral lifestyle and pathogenicity frequency

We used bootstrapping to assess the significance of observed changes in viral lifestyle ratios and pathogenicity gene content between sample categories (**Figures 3a and 4a**). To do this, we generated sample distributions, medians, and 95 % CIs of the medians over 1000 bootstrapping iterations with replacement. In these analyses, non-overlapping 95 % CIs show that significant differences exist between sample categories (p < 0.05; code available at https://github.com/hopefulmonstersucla/coprolite_viromes).

### Accommodating sparse pre-modern samples via down-sampling

Finally, given the low sample size of the pre-modern cohort, we implemented a down-sampling analysis to determine if significant signal between pre-modern and modern samples could still be detected if the modern samples were down-sampled to the size of the pre-historic cohort. This was done on the viral lifestyle ratios and virulence factor prevalence analyses (**Figures 3a** and **4a**). For each metric, modern samples were repeatedly resampled with replacement to match the number of pre-modern samples, and summary statistics were recalculated for each iteration. Specifically, median values were computed across resampled datasets, generating empirical distributions of medians that reflect uncertainty due to sample size.

## Results

### Samples analyzed

To investigate how viral communities in the human gut have changed over time, we extracted phage contigs and genomes from 14 publicly available pre-modern coprolite microbiome samples from sites in modern-day Austria, n = 3; Mexico and United States, n = 8; and South Africa, n = 3; **Figure 1a; Supplementary Table 1**). These were complemented by 492 samples from industrialized populations (China, n = 74; Denmark, n = 109; Spain, n = 140; USA, n = 169; **Figure 1a**) and 502 samples from non-industrialized populations (Fiji, n = 174; Madagascar, n =112; Mexico, n = 22; Nepal, n = 56; Peru, n = 36; Tanzania, n = 102; **Figure 1a**). Pre-modern samples spanned from approximately 1301 BCE to 1400s CE, and all non-industrialized and industrialized samples were collected from contemporary, live people (**Figure 1b**). Samples were from agrarian and hunter-gatherer societies, and from both rural and (sub)urban populations (**Figure 1c**).

### Bacteriophage functional genomic relatedness over time

To examine the functional relatedness between bacterial and archaeal viruses (phages) from pre-modern, non-industrialized, and industrialized population samples, we generated viral metagenomically-assembled genomes and a protein-sharing network (APGs; **Figure 2a**). This yielded n = 42, n = 2316, n = 903 pre-modern, non-industrialized, and industrialized APGs, respectively, which we then analyzed in conjunction with n = 4534 database genomes^43^.

We then used the network to determine whether observed protein-sharing frequencies differed from random expectation by comparing observed values to null distributions generated by permutation (p < 0.05; grey 95 % CI shown; **Figure 2b**). Observed frequencies (red lines) above the null distribution indicate greater-than-expected genomic similarity, whereas frequencies below the null distribution indicate reduced similarity.

This analysis showed that pre-modern genomes (PM) did not cluster with themselves, with observed connection frequencies consistent with random expectation (observed = 0.000017, 95 % CI_null_ = 0.000000 - 0.000316; **Figure 2b**). In contrast, non-industrialized (NI), industrialized (I), and database (DB) genomes all clustered significantly within group (observed = 0.203549, 0.045476, and 0.537590 versus 95 % CI_null_ = 0.120222 - 0.128900, 0.025271 - 0.030885, and 0.521334 - 0.554589 for NI-NI, I-I, and DB-DB comparisons, respectively). Pre-modern genomes also clustered away from non-industrialized, industrialized, and database genomes (observed = 0.002946, 0.001042, and 0.000848 versus 95 % CI_null_ = 0.004962 - 0.008193, 0.001731 - 0.004234, and 0.007895 - 0.010113 for PM-NI, PM-I, and PM-DB comparisons, respectively). While contemporary genomes exhibited greater within-group similarity than expected by chance, connectivity between non-industrialized and industrialized genomes was consistent with random expectation (observed = 0.118901, 95 % CI_null_ = 0.112262 - 0.123404). Similar to the pre-modern APGs, both contemporary groups clustered away from database genomes (**Figure 2b**), highlighting the novelty of the gut virome relative to the broader landscape of currently characterized viral genomic diversity (**Figure 2a**).

### Changes in Viral Lifestyle over Time

Given the observed changes in gut virome gene-content similarity, we next investigated whether the gut viruses had also changed lifestyle over time. To investigate this, we identified temperate and lytic phage contigs in our metagenomic contigs using PhaTYP on contigs conservatively labeled as phage using PhaMER^39,46^ (**Supplementary Figure 1**). Doing so showed that the temperate-to-lytic ratio (TLR) of phage contigs significantly increased over time and industrialization, with the highest values seen in industrialized samples (**Figure 3a**; pre-modern TLR, 95 % CI: 1.16, 0.91 - 1.28; non-industrialized TLR: 1.33, 1.32 - 1.34; industrialized TLR: 1.37, 1.35 - 1.38; bootstrapped median, 95 % CIs over 1000 iterations). At the population/country level, pre-modern samples spanned both lytic- and temperate-dominated viromes (TLR range = 0.78 - 1.53), whereas all contemporary populations were temperate dominated (TLR range = 1.25 - 1.49; **Figure 3a**). We then confirmed that this outcome was not a result of unbalanced replication between pre-modern and contemporary datasets by downsampling contemporary samples to the size of the pre-modern cohort. The same ordering was retained following downsampling to accommodate the unbalanced sample numbers in pre-modern vs modern samples sets (non-industrialized TLR: 1.33, 1.25 - 1.41; industrialized TLR: 1.37, 1.28 - 1.45), indicating that the increase in temperateness was robust to differences in sample size (**Figure 3a**). Together, pooled, downsampled, and population-level analyses support a shift toward increasingly temperate gut viromes in contemporary human populations.

We independently validated this pattern using assembled phage genomes (APGs) and CheckV-derived lifestyle annotations (**Figure 2a**, **Figure 3b**)^40^.

Consistent with the metagenomic analysis, genomic temperate-to-lytic ratios increased from 0.14 to 0.26 to 0.31 across pre-modern, non-industrialized, and industrialized populations, respectively (**Figure 3b**). These values corresponded to temperate genomes comprising 11.9 %, 20.5 %, and 23.4 % of recovered APGs (n = 42, n = 2316, n = 903 APGs, respectively). Together, these metagenomic and genomic analyses provide independent lines of evidence that the human gut virome has become increasingly temperate over time and with industrialization.

### Increased prevalence of pathogenicity genes in contemporary phages

Given the association between temperate viral lifestyles and the phage-mediated transfer of genes that can modulate bacterial pathogenicity (i.e., virulence factors)^25,47^, we next examined the prevalence and composition of these genes as a proxy for how gut viruses might influence bacterial pathogenicity. In contrast to viral lifestyles, which shifted primarily between pre-modern and contemporary populations, virulence factors showed a pattern more closely associated with industrialization (**Figure 4a**). Across pooled populations, the proportion of phage proteins annotated as virulence factors was similar between pre-modern and non-industrialized samples (0.118, 95% CI = 0.106-0.128 and 0.118-0.119, respectively) but increased in industrialized populations (0.123, 95% CI = 0.122-0.124). Downsampling to account for the smaller pre-modern sample size recovered the same pattern (PM = 0.118, 95% CI = 0.106-0.128; NI = 0.118, 95% CI = 0.113-0.124; I = 0.123, 95% CI = 0.117-0.130).

Population-level analyses likewise supported this trend, with contemporary populations generally occupying the upper end of the observed range (**Figure 4a**). Together, pooled, downsampled, and population-level analyses indicate that phage-associated virulence factors are enriched in industrialized gut viromes. Among the 100 most differentially represented virulence factors, genes associated with adherence, immune modulation, and regulation were the dominant functional categories, accounting for 27, 16, and 13 factors, respectively (**Figure 4b**), with the remaining factors distributed across a range of less common functional classes (**Supplementary Tables 3 and 4**).

## Discussion

Human diets, lifestyles, and environments have changed from pre-modern to modern times, driving changes in the human gut microbiome and declines in enteric human health^3,15–17^. Although the short-term establishment and evolution of the gut virome have been examined^8,9,18,21^, little is known about how the viral community in the human gut has changed across pre-modern to industrialized eras^7^. To address this, we aggregated and compared fecal metagenomic samples from current-day industrialized^6,32^ and non-industrialized populations^6,33^, as well as from pre-modern coprolites spanning from 1301 BCE – 1400s CE^6,30,31^.

This allowed us to directly characterize the trajectory of the human gut virome over time and with industrialization. While acknowledging the well-documented caveats of using ancient DNA (aDNA) approaches, including DNA degradation, preservation bias, and limited sample availability^27,28^, our conservative metagenomic and genomic analyses provide direct evidence that human gut viral communities have become (i) more similar in gene content, (ii) more temperate, and (iii) increasingly supportive of bacterial pathogenicity.

While there is an emerging association between altered bacterial physiology, enhanced temperate infection, and ecosystem dysbiosis in other environments^22–25,49,50^, the ecological and health implications of shifts in viral lifestyles on the function of the human gut microbiome are unclear. However, prior research in other ecosystems shows that a shift toward temperateness leads to suppressed lytic control and subsequent proliferation of copiotrophic bacteria^23,25,51^. In the gut, reduced lytic infection likely alters community composition^5,11,13^. Increased pathogenicity gene prevalence in gut phage genomes could further enhance bacterial activity associated with gut inflammation, autoimmunity, and associated diseases in industrialized societies with temperate viromes^19,26,52–54^. Altogether, it is likely that the pre-modern gut had higher lytic viral titers, lower bacterial densities, and a virome and microbiome that had a less antagonistic relationship with their human habitat.

There have been wholesale changes in human society since the pre-modern era^16,17,29,55^. Any combination of these changes may underpin observed shifts in viral lifestyle, pathogenicity, and function. In particular, modern dietary changes that enrich the gut in simple sugars may have enhanced bacterial growth rates, promoted a switch from lytic to temperate viral infection, and especially in industrialized populations, elevated bacterial pathogenicity and immunogenicity^3,15^. It remains unclear whether the human gut microbiome is dysbiotic or if it has reached a new homeostasis with current conditions^5,13^. It is also uncertain whether restoring the gut microbiome to a pre-modern state would be beneficial without concurrent changes in living conditions, diet, and other factors. However, where microbiome modification could benefit health, our findings suggest that gut viral communities may toggle between a high-resource, temperate, pathogenic state and a more lytic, health-associated state. This presents a potential ecological lever to reinvigorate lytic infection (“gut reviralization”) as part of a dietary therapeutic approach grounded in the long-term ecology of the gut.

## Code and Data Availability

Bioinformatic code and Assembled Phage Genomes (APGs) generated in this study can be found at https://github.com/hopefulmonstersucla/coprolite_viromes. This code can be used from SRA download of all samples to our analytical pipeline (**Supplementary Figure 1**).

## Acknowledgements

First, The authors would like to thank the two anonymous reviewers for their insightful and constructive comments, which greatly strengthened the rigor, clarity, and overall quality of this manuscript. The authors would also like to express their gratitude to Alex Hoffmann and the UCLA Institute for Quantitative and Computational Biosciences for their support. This work was supported by NSF Award BIO-OCE 2201645 to BK, which also provided stipends to AK, ML, JK, SB, NF, MK, BP, and IT to attend the UCLA Institute for Quantitative and Computational Biosciences B.I.G. Summer program. This work used computational and storage services associated with the Hoffman2 Shared Cluster provided by UCLA Office of Advanced Research Computing’s Research Technology Group.

## Supplementary Tables and Figures

**Supplementary Table 1:** Summary of samples For each sample, general information about a sample’s origin, sequenced reads, and assembly are provided.

<u>Supp_table_1.xlsx</u>

https://docs.google.com/spreadsheets/d/1OIpQhEROnfcVXjUH9Vd1klP6bOpYc4K1/edit?gid=419351271#gid=419351271

**Supplementary Table 2: L**ytic-to-temperate ratio of all samples Summaries of the number of lytic and temperate phage contigs per sample.

<u>Supp_table_2.xlxs</u>

https://docs.google.com/spreadsheets/d/11jE2biKnLu6mMxHNCBI3l53xGL6c_0D_fl1Xq9a64Gg/edit?usp=drive_link

**Supplementary Table 3:** The names of Virulence Factors in **Figure 4b**

**Figure 4b** contains the top 100 most differentially represented virulence factors across samples. **Supplementary Table 3** also contains the VF name, category, and functional description from the VFDB. **Supplementary Table 3** also contains the p-value of each factor across all samples.

<u>supp_table_3.xlsx</u>

https://docs.google.com/spreadsheets/d/1Th6hosTLvCiz2tQ84EPEIZQvjOwSA8T4/edit?gid=278719906#gid=278719906

**Supplementary Table 4:** Differentially represented VFC counts

Based on the VFs in **Figure 4b** and **Supplementary Table 3**, **Supplementary Table 4** provides the raw counts of how each factor in the top 100 most differential represented virulence factors is categorized by the VFDB.

<u>Supp_table_4.xlsx</u>

https://docs.google.com/spreadsheets/d/1xz_qHkDIoEXwhWnMLmaeVXAzIJ8LjaT7RWFWh8LWjBw/edit?usp=drive_link

**Supplementary Figure 1:**
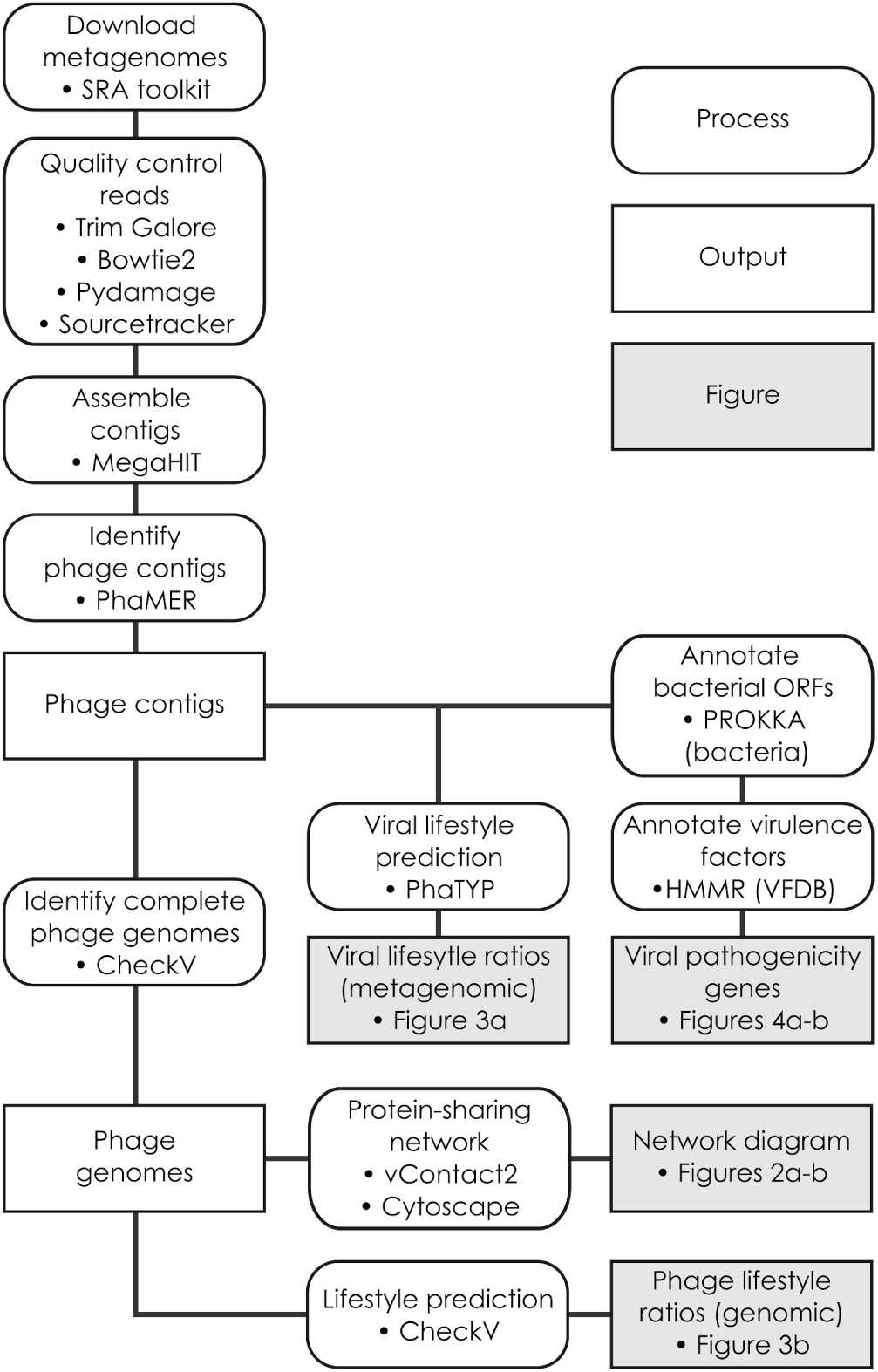
Bioinformatic Pipeline including sample aggregation and processing, and metagenomic and genomic analyses. This includes sample downloading (provenance shown in Figure 1), quality control (removal of low quality reads, removal of human contamination, retention of only reads with hallmarks of ancient providence, and removal of non-human-associated samples using TrimGalore, Bowtie2, Pydamage, and Sourcetracker, respectively), assembly into contigs, and extraction of phage contigs to make the ‘Phage contigs’ pool for the metagenomic analyses of viral lifestyles and viral pathogenicity genes shown in Figures 3 **and 4**, respectively. Complete phage genomes were also extracted from the Phage contigs pool using CheckV and used to populate a protein-sharing network using vContact2, assessed for lifestyle using CheckV outputs, with analyses shown in Figures 2 **and 3**, respectively. See Methods for further details.

## Notes

### Competing Interest Statement

The authors have declared no competing interest.

### Summary of Updates

This revision reflects accommodation of highly constructive reviewer comments that especially allowed the increased rigour of bioinformatic quality control.

